# Tetraploid *Caenorhabditis elegans* embryos exhibit enhanced tolerance to osmotic stress

**DOI:** 10.1101/2025.10.13.678412

**Authors:** Yuiko Oyama, Misako Ohama, Namu Yamada, Nozomi Suzuki, Kenji Kimura, Yuki Hara

## Abstract

Polyploidy is a widespread phenomenon in development and evolution and is often associated with altered cellular physiology and developmental robustness. However, the influence of genome doubling on embryonic robustness to environmental stress remains poorly understood. Here, we investigated developmental traits and osmotic responses in tetraploid *Caenorhabditis elegans* embryos. Tetraploid animals exhibited enlarged body and tissue sizes and produced embryos with approximately 1.6-fold greater volume than diploids, accompanied by moderately delayed development and partial embryonic arrest. When exposed to a range of osmotic environments, both diploid and tetraploid embryos swelled or shrank in response to external osmolarity. Strikingly, tetraploid embryos at the early stage maintained normal cell division across a broader range of osmotic conditions than diploids. Quantitative analyses further revealed that tetraploid embryos exhibit reduced cytoplasmic mass density, largely reflecting lower protein concentration, while lipid and RNA levels remain unaffected. These compositional differences likely buffer volume fluctuations and underlie the enhanced osmotic tolerance of tetraploid embryos. Our findings demonstrate that genome doubling reshapes embryonic cellular physiology in a non-proportional manner to ploidy, thereby enhancing robustness during early development.

## INTRODUCTION

Polyploidy, the presence of more than two sets of chromosomes, is a common feature of metazoan development. Polyploid cells arise in diverse tissues, including brain, liver, bone marrow, muscle, and placenta in diploid organisms (Sagi and Benvenisty, 2017), and can modify cellular properties to confer specialized physiological functions. A well-known consequence of cellular polyploidization is an increase in cell size, observed both across cell types within an organism and among eukaryotic species (Gillooly et al., 2015; Gregory, 2001). Larger cells support the formation of larger tissues and accelerate tissue repair processes following injury (Bailey et al., 2021; Fox et al., 2020; Lessenger et al., 2025). In addition to increased cell size, polyploidization induces multiple alterations in the transcriptome and organelle morphology. For instance, nuclear size scales with DNA content across species and adjusts dynamically when DNA content is experimentally manipulated (Heijo et al., 2020; Heijo et al., 2022). Although transcript abundance generally scales with gene copy number according to ploidy, recent transcriptomic analyses have revealed that polyploidization does not lead to proportional increases in gene expression (Lanz et al., 2022; Lessenger et al., 2025; Yahya et al., 2022). Similarly, metabolic scaling in polyploid *Xenopus laevis* deviates from ploidy proportionality (Cadart *et al*., 2023). These findings suggest that polyploidization alters the relative abundance of various molecular species within the cell in a non-proportional manner. Therefore, cells must possess physiological mechanisms that adjust ploidy-dependent alterations in cellular properties to maintain homeostasis under environmental and physiological stresses. Indeed, whole-body genome duplication can impair development and disrupt tissue organization in some species, including mice and zebrafish, where induced polyploidy leads to developmental arrest (Imai *et al*., 2020; Small *et al*., 2021). By contrast, polyploid embryos are viable in several species, including plants, reptiles, amphibians, and nematodes (Cadart et al., 2023; Clarke et al., 2018; Fankhauser, 1945; Robinson et al., 2018). These observations indicate that the physiological consequences of polyploidization vary among species and that genome doubling can reshape multiple aspects of cellular physiology in a manner that is not simply proportional to ploidy. In particular, during embryogenesis, when cellular properties undergo rapid reorganization, such alterations could have a profound influence on developmental robustness. However, the influence of genome doubling on embryonic physiology and stress tolerance remains poorly understood. Here, we examined how polyploidization affects developmental traits and osmotic responses in tetraploid *Caenorhabditis elegans* embryos.

## RESULTS AND DISCUSSION

### Tetraploid worms exhibit reduced embryonic viability and reproductive capacity

To assess the developmental impact of tetraploidy in *C. elegans*, we first compared adult body length between diploid and tetraploid strains. Two tetraploid strains were used: SP346, derived from the wild-type diploid N2 strain, and ECB003, newly generated in this study from the diploid NF4496 strain expressing fluorescently tagged proteins. Although occasional reversion to diploid state was detected, both tetraploid strains were relatively stable in ploidy, consistent with previous reports (Clarke *et al*., 2018). In aged tetraploid worms, oocytes carrying haploid genomes were rarely detected in the gonad (Fig. S1A).

Based on the current understanding of oogenesis and meiosis (Hillers *et al*., 2017), such ploidy reduction likely occurs during mitotic divisions of germline stem cells within the mitotic zone of the gonad. Age-dependent alterations in intracellular properties, including nuclear structure (D’Angelo et al., 2009), may also contribute to reduced mitotic fidelity in germline stem cells. Therefore, relatively young adult worms were used for analyses in this study.

Tetraploid ECB003 worms were approximately 1.5-fold longer than the diploid adults (Fig. 1A) as previously reported in other tetraploid strains (Clarke et al., 2018; Misare et al., 2023). This increase in body size extended to the reproductive tissues, with both gonads and embryos substantially enlarged in tetraploid worms (Fig. 1A). The ECB003 strain, which expresses fluorescently tagged histone and a plasma membrane marker, allowed direct visualization of chromatin and plasma membrane during oogenesis in the gonad. In *C. elegans* gonads, germ cells grow by acquiring cytoplasmic components through intercellular bridges before undergoing cellularization to form individual oocytes (Wolke et al., 2007) (Fig. 1A, the diploid gonad in NF4496). In the tetraploid gonad, numerous immature germ cells were observed even beyond the loop region, and cellularization appeared markedly delayed (Fig. 1A, ECB003). Notably, the number of cellularized oocytes was reduced compared with diploids (Fig. 1A), consistent with previous observation (Misare *et al*., 2023). Since germ cell morphology and oocyte formation are regulated by actomyosin-dependent cortical tension and intercellular bridge closure (Kimura and Motegi, 2025), these processes may be partially impaired in tetraploids.

**Figure 1.**
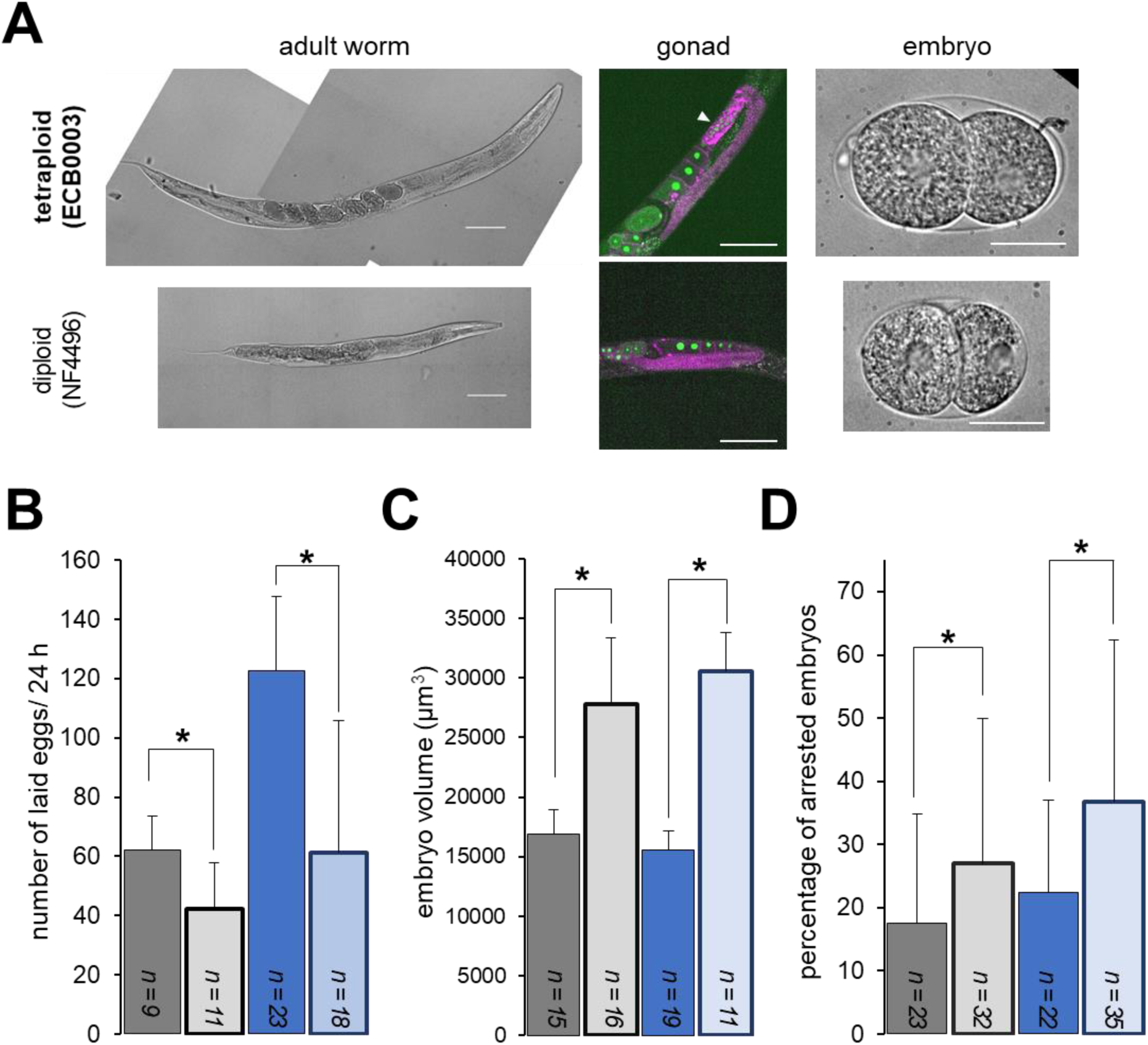
Developmental and morphological profiles of tetraploid worms. (**A**) Representative images of adult worms (DIC, scale bar: 100 µm), gonad region (magenta, mCherry PH-domain; green, GFP histone H2B; scale bar: 100 µm), and 2-cell-stage embryos (DIC, scale bar: 20 µm) from tetraploid strain ECB003 and diploid strain NF4496. Arrowhead indicates the gonadal region with labelled PH-domain accumulation. (**B**) Mean numbers (± s.d.) of eggs laid from an adult worm for 24 h. (**C**) Mean embryo volume (± s.d.) at 2-cell stage. (**D**) Percentage (± s.d.) of embryos that arrested before reaching the larval stage (without hatching from eggshell) within 48 h after egg laying. Asterisks indicate significant differences between tetraploid and diploid embryos (Student’s t-test, *P* < 0.05).

Reproductive output was also impaired in the tetraploid. Brood size, measured as the number of eggs laid within 24 hours, was significantly smaller in tetraploids than in diploids (Fig. 1B), consistent with earlier reports (Clarke *et al*., 2018; Misare *et al*., 2023). Reduced egg production was a general characteristic of tetraploids, though the extent varied between strains. Following fertilization, tetraploid embryos were significantly larger than diploids, both at the whole-embryo level (Fig. 1C) and at the individual blastomere level (Fig. S1B, C), showing an approximately 1.6-fold increase in volume across strains, consistent with previous studies (Clarke *et al*., 2018; Misare *et al*., 2023). Embryonic lethality, although modest, was significantly higher in tetraploids than in diploids, whether assessed on glass slides (Fig. 1D) or on growth plates (Fig. S1D). Most lethal embryos progressed through multiple rounds of cell divisions, often exceeding the 100-cell stage, where individual blastomeres became indistinct under a dissecting microscope. As these stages occur after zygotic genome activation (Powell-Coffman *et al*., 1996), the observed lethality likely reflects defects in later embryogenesis rather than early developmental events. Furthermore, developmental progression was generally slower in tetraploid embryos, suggesting that reduced mitotic fidelity—such as chromosome segregation errors and impaired cell cycle checkpoint regulation—contributes to diminished embryonic viability.

### Higher tolerance to osmotic stress in tetraploid embryos

Tetraploid embryos are prone to developmental arrest, possibly due to chromosome segregation errors during cell division, yet they produce larger eggs. The biological significance of this phenomenon remains unclear. A previous study (Misare et al., 2023) demonstrated that tetraploid larvae exhibit increased resistance to drug treatments. To evaluate whether tetraploid embryos differ in stress tolerance, we exposed embryos to a range of osmotic conditions using conventional egg salts (ES) buffer: standard (100% ES), diluted (20–90% ES), and concentrated (110–200% ES) conditions (Fig. 2A). Under extreme hypoosmotic (20% ES) or hyperosmotic (200% ES) conditions, both diploid and tetraploid embryos swelled or shrank mildly, respectively. Thus, the eggshell of tetraploids retained its barrier function, maintaining high embryonic viability across external osmotic conditions. To assess the osmotic sensitivity independent of eggshell, we performed RNAi knockdown of *perm-1*, a gene required for eggshell impermeability (Carvalho *et al*., 2011). Under hyperosmotic conditions (200% ES), both diploid and tetraploid *perm-1(RNAi)* embryos exhibited excessive shrinkage, generating large cavities between the embryo and the eggshell (Fig. 2A). Conversely, under hypoosmotic conditions (20% ES), embryos swelled markedly, with no visible cavity between the embryo and the eggshell (Fig. 2A).

**Figure 2.**
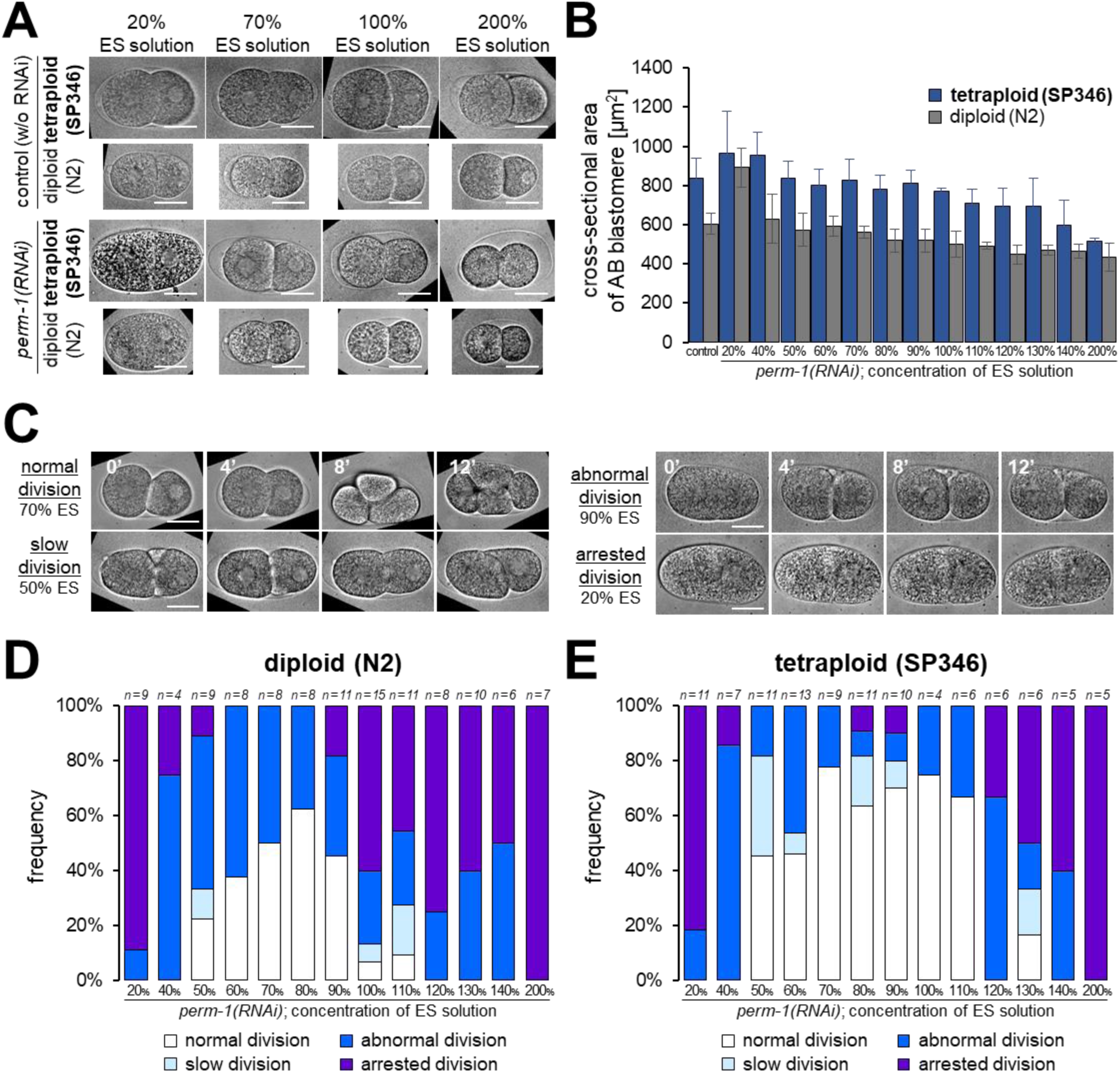
Osmotic response in tetraploid embryos. (**A**) Representative images of diploid (N2) or tetraploid (SP346) embryos, with or without *perm-1(RNAi)*, after exposure to different concentrations of egg salt (ES) solution. Scale bar: 10 µm. (**B**) Mean cross-sectional area (± s.d.) of AB blastomeres in diploid (N2) and tetraploid (SP346) *perm-1(RNAi)* embryos incubated in different ES concentrations. (**C**) Representative snapshots of categorized cell division phenotypes in *perm-1(RNAi)* embryos after exposure to varying ES concentrations. Scale bar: 10 µm. (**D, E**) Frequencies of categorized cell division phenotypes in diploid (**D**, N2) and tetraploid (**E**, SP346) *perm-1(RNAi)* embryos across ES concentrations. Numbers of embryos analyzed (*n*) are indicated.

Measurements of whole-embryo and AB blastomere sizes across ES concentrations confirmed concentration-dependent swelling under hypoosmotic conditions (20–40% ES) and shrinkage under hyperosmotic conditions (>110% ES) in both diploid and tetraploid embryos (Fig. 2C, D). Similar trends were observed in P1 blastomeres (Fig. S2B) and in different genetic backgrounds (Fig. S2C, D). Interestingly, the degree of swelling in tetraploid embryos under hypoosmotic conditions was less pronounced than the shrinkage observed under hyperosmotic conditions (Fig. S2A).

Despite these morphological changes, many *perm-1(RNAi)* embryos displayed cell division defects under both hypo- and hyperosmotic conditions (Fig. 2B). Moderate phenotypes included delayed cell cycle progression, abnormal nuclei (e.g., multiple nuclei within individual blastomeres), and blastomeres lacking DNA, likely resulting from chromosome segregation errors. Severe phenotypes involved complete developmental arrest, with nuclei and spindles remaining in a static intracellular position, indicative of disrupted intracellular homeostasis. To systematically evaluate these phenotypes, we classified the defects into mild, moderate, and severe categories and quantified their frequencies across ES concentrations (Fig. 2E, F). In the diploid N2 strain, normal cell division occurred predominantly between 50–110% ES, peaking around 80% ES, with defects increasing as osmotic conditions deviated from this optimum (Fig. 2E). Tetraploid SP346 embryos showed a similar peak concentration (Fig. 2F). However, the range of ES concentrations supporting normal cell division was broader, as reflected by a wider Gaussian distribution (Fig. S2E, F). While the optimal ES concentration was comparable between diploid and tetraploid embryos, the overall probability of normal development was slightly higher in tetraploids. Consistent trends were obtained in the other genetic backgrounds (NF4496 and ECB003), although their overall frequencies of normal division, including delayed division, were lower than in the N2 and SP346 strains for reasons yet unclear (Fig. S2G, H). Collectively, these results demonstrate that tetraploid embryos exhibit enhanced tolerance to osmotic variation, suggesting enhanced robustness against environmental stress, likely mediated by altered cellular physiology and internal osmotic regulation.

### Differences in cellular composition between tetraploid and diploid embryos

Cellular responses to osmotic stress are largely determined by the intracellular concentration of molecules and organelles, which act as osmolytes given the restricted permeability of the plasma membrane (Wu *et al*., 2022). These molecular profiles are dynamically regulated throughout the cell cycle and development, and also vary spatially within individual cells. This raises the possibility that polyploidization alters intracellular composition in embryos. As a first step, we measured the refractive index (RI) of whole embryos by holotomography (Fig. 3A), which reflects cellular mass density, with higher RI values corresponding to higher molecular density (Biswas et al., 2025; Kim and Guck, 2020). Since RI values gradually increase during embryogenesis (Fig. S3A), we restricted our analyses to early embryonic stages (≤28-cell stage). Both diploid and tetraploid embryos exhibited heterogeneous distribution of RI values, with nuclei displaying lower values, consistent with previous analyses in *C. elegans* embryos (Biswas *et al*., 2025; Goda *et al*., 2024).

**Figure 3.**
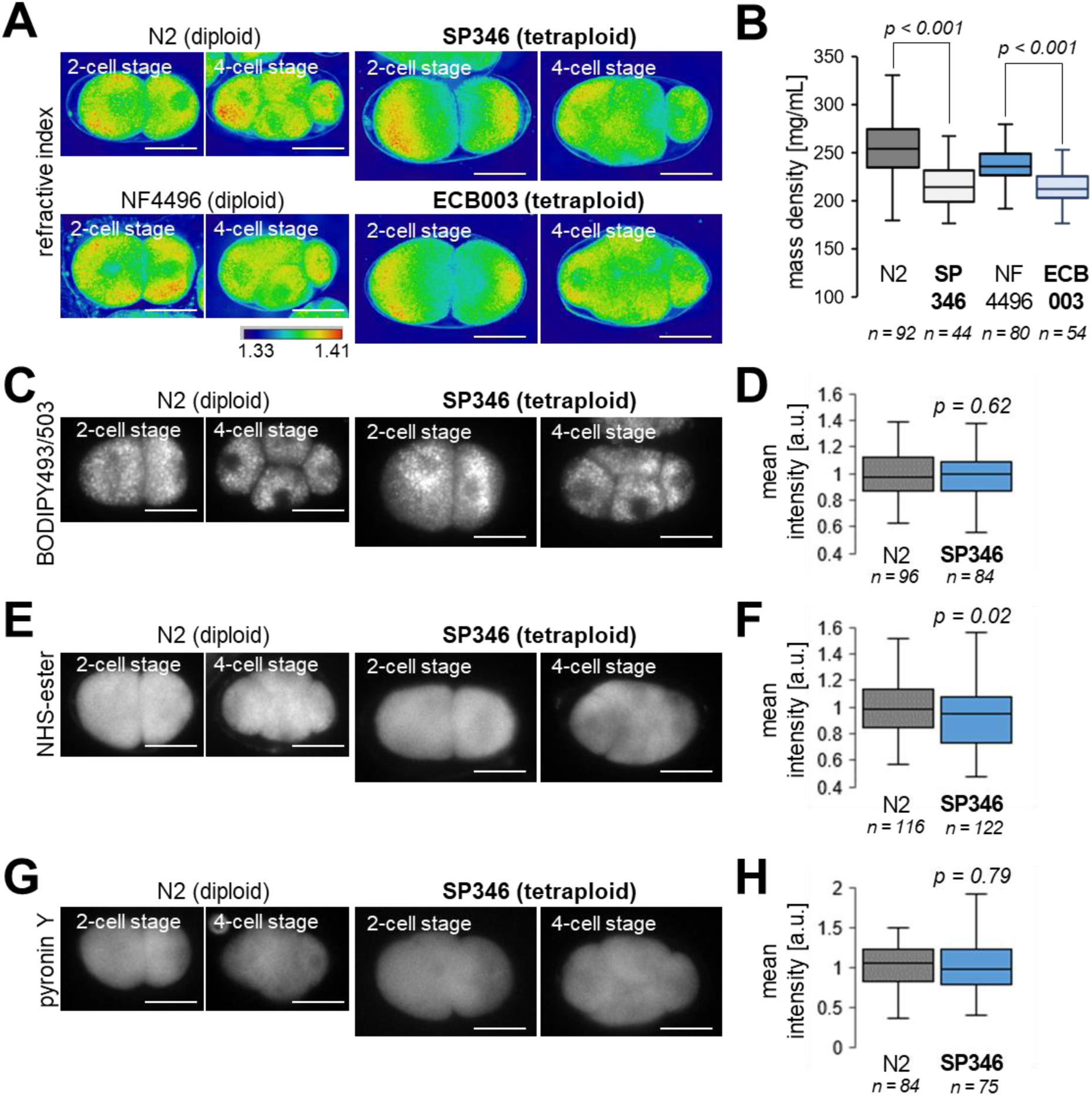
Cytoplasmic composition in tetraploid embryos. (**A**) Representative refractive index (RI) distribution of diploid (N2, NF4496) and tetraploid (SP346, ECB003) embryos at 2- or 4-cell stages. Color scale indicates relative RI values. Scale bar: 10 µm. (**B**) Box plots of calculated cellular mass density derived from RI imaging. The line within the box represents the median. The upper/lower lines represent the 75th/25th percentiles. The whiskers represent the 95th/5th percentiles. *p*-values are from Wilcoxon tests comparing diploid and tetraploid embryos; numbers of embryos analyzed (*n*) are indicated. (**C, E, G**) Representative fluorescence images of diploid (N2) and tetraploid (SP346) embryos at 2- or 4-cell stages stained with BODIPY 493/503 (lipids, **C**), NHS-ester (proteins, **E**), and pyronin Y (RNAs, **G**). Scale bar: 10 µm. (**D, F, H**) Box plots showing normalized fluorescence intensities of BODIPY 493/503 (**D**), NHS-ester (**F**), and pyronin Y (**H**). Intensities were normalized to mean diploid values.

Furthermore, RI values were relatively higher around predicted centrosome position, likely reflecting local enrichments of yolk granules and organelles transported by microtubule-based motor proteins (Kimura and Kimura, 2011). Importantly, the mean mass density of entire embryos, estimated from RI imaging, was approximately 10% lower in tetraploid embryos than in diploid (Fig. 3B). Because intracellular components are limited in tetraploid embryos, their uneven distribution within the cytoplasm may accentuate relatively low-density regions, such as the side opposite to centrosomes (Fig. 3A). Notably, our calculated mass densities obtained by RI measurement through holotomography were almost consistent with previously reported values from precise measurements in *C. elegans* diploid embryos (Biswas *et al*., 2025). Together, these results suggest that intracellular components do not scale with cell volume in tetraploid embryos, resulting in reduced mass density that may buffer swelling under hypoosmotic stress.

To search for the molecular species contributing to the mass density reduction, we quantified representative biomolecules—lipids, proteins, and RNAs—using specific fluorescent dyes. Neutral lipids, visualized by BODIPY 493/503, appeared as cytoplasmic puncta (Fig. 3C), likely corresponding to visible yolk granules in DIC images (Fig. 2A) or fluorescently labelled yolk proteins reported previously (Lessenger *et al*., 2025), and dot-like high-density signals in RI imaging (Fig. 3A). Despite such biased distribution, mean BODIPY intensity was comparable between diploid and tetraploid embryos (Fig. 3D). This indicates that lipid concentration and yolk granule distribution are similar between ploidy states, although their molecular composition, including vitellogenin proteins, may differ. Next, overall protein abundance, assessed by NHS-ester staining of peptide bonds, was approximately 10% lower in tetraploid embryos compared with diploid embryos, with a slightly higher intensity near centrosomes (Fig. 3E, F). Given their ∼1.6-fold greater volume, this implies ∼1.4-fold more total protein content in tetraploid embryos, but a dilution in concentration that closely parallels RI measurement, suggesting that proteins are the primary contributors to the reduced mass density. In contrast, total RNA levels, measured using pyronin Y, revealed no significant difference between diploid and tetraploid embryos (Fig. 3G, H). Together, these results suggest that proteins, rather than lipids or RNAs, underlie the reduced mass density in tetraploid embryos.

Since acute increases in cell volume coincide with dilution of protein concentration in some eukaryotic species (Neurohr et al., 2019), we asked whether this protein dilution simply reflects enlarged embryo size in tetraploids. To test this, we examined enlarged diploid embryos generated by *C27D9.1(RNAi)* (Hara and Kimura, 2009; Weber and Brangwynne, 2015). Despite having comparable volumes to tetraploid embryos (Fig. S3B, C), these embryos displayed unaltered lipid, protein, and RNA concentrations (Fig. S3D–F), consistent with prior evidence that maternally loaded oocyte components (e.g., nucleolar materials) scale independently of cell size (Weber and Brangwynne, 2015). Thus, reduced protein concentration in tetraploid embryos is not a simple consequence of larger embryonic size. Instead, it may reflect impaired molecular composition of the gonadal cytoplasm supplied into oocytes during oogenesis, consistent with the abnormal gonadal architecture observed in tetraploid worms (Fig. 1A). Programmed intestinal polyploidization generally contributes to boost yolk production and supply it from intestinal cells to oocytes (Lessenger *et al*., 2025). Thus, excessive genome doubling in tetraploid tissues may disrupt balanced biosynthetic output, leading to dysregulated biosynthesis and imbalanced molecular composition in oocytes. This protein dilution may have broad physiological consequences. Ribosome concentration is positively correlated with growth rate in *C. elegans* (Uppaluri *et al*., 2016), and reduced ribosomal protein abundance may underlie the slower development and shorter lifespan of tetraploid (Misare *et al*., 2023). Similarly, experimentally induced dilution of cytoplasmic composition in some eukaryotic cells induces cellular senescence with cell cycle arrest (Neurohr et al., 2019). Furthermore, the observed ∼1.4-fold increase in the total protein content, which is less than the expected 2-fold relative to DNA content, implies altered DNA-to-cellular molecule ratios. DNA-to-cytoplasm ratios are long recognized as critical regulators of early embryonic events, including cell cycle progression, cellular motility, and zygotic transcription (Amodeo et al., 2015; Newport and Kirschner, 1982a; Newport and Kirschner, 1982b). Imbalanced DNA-to-cytoplasmic molecule ratios in tetraploids may therefore perturb nuclear and cytoplasmic activities, contributing to slow growth and developmental abnormalities. Future transcriptomic and proteomic analyses will be essential to identify selectively regulated pathways and to clarify how ploidy-dependent molecular scaling confers robustness to osmotic stress.

## MATERIAL AND METHODS

### Worm strains

*C. elegans* strains used in this study include N2 (wild-type), SP346 (N2-derived tetraploid), and NF4496 [expressing GFP::Histone H2B (*his-11*) and mCherry::PH(PLCdelta-1) by mating EG4601 and OD70 strains, kindly distributed from Dr. Kiyoji Nishiwaki]. All strains were obtained from the *Caenohabditis* Genetic Center (CGC). The tetraploid derivatives of NF4496, designated as ECB003, were generated following the established methods (Clarke *et al*., 2018). Briefly, L3-L4 stage diploid worms were subjected to feeding RNA-mediated interference (RNAi) targeting the *rec-8* gene. F2 progeny were screened under a stereo microscope, and individuals exhibiting longer body length to diploid controls were selected. These candidate worms were cultured individually, and their ploidy status was verified by staining whole worm with Hoechst 33342 to assess oocyte DNA content in the gonad. Since the tetraploid strains often revert to diploid, longer adult worms were selectively transferred to fresh nematode growth medium (NGM) agar plates [51.3 mM NaCl, 0.25%(*w/v*) Bacto Peptone, 1.7%(*w/v*) Bacto Agar, 5 µg/ml cholesterol; 1 mM CaCl_2_; 1 mM MgSO_4_; 25 mM KH_2_PO_4_ (pH 6.0)] for maintaining the tetraploid status.

### RNA mediated interference

RNAi by feeding was performed for *rec-8, perm-1,* and *C27D9.1. E. coli* carrying the corresponding gene sequences in the L4440 vector (in both orientations) were obtained from the Ahringer RNAi laboratory ((Kamath *et al*., 2003); supplied by Danaform Japan). Single colonies were inoculated into 2 mL of Luria Broth (LB) media containing 100 µg/mL ampicillin and grown overnight. For standard RNAi induction (excluding for *perm-1*), 1:100 (v/v) of the pre-culture was inoculated into fresh LB media containing 100 µg/mL ampicillin and incubated at 37°C for 4 h.

Subsequently, 1 mM of isopropyl β-D-thiogalactopyranoside (IPTG) was added, and cultures were further incubated for 1 hour to induce dsRNA expression. For each RNAi plate, 200 µL of the induced culture was spread onto 60 mm NGM agar plates containing 1 mM IPTG and 100 µg/mL ampicillin. RNAi plates were incubated at 25°C for 14 hours, then stored at 4°C until use. For *perm-1* RNAi, pre-cultures of *E. coli* carrying *perm-1* sequence in L4440 vector or the empty control L4440 vector was mixed at a 1:1 ratio before inoculation. 200 µL of induced culture was spread onto 60 mm NGM agar plate containing 10 µM IPTG and 100 µg/mL ampicillin. For RNAi treatment, 20–30 L2–L3-stage larvae were transferred to the prepared RNAi plates, and incubated for 24–48 h (for *perm-1*) or 48 h (for *rec-8, C27D9.1*) prior to imaging.

### Live imaging of adult worms and embryos

Adult wild-type *C. elegans* were dissected into M9 buffer (41.8 mM Na_2_HPO_4_; 22 mM KH_2_PO_4_; 85.5 mM NaCl; 1 mM MgSO_4_) directly on a glass slide. Released embryos were transferred into M9 buffer on an eight-well glass slide (TF0808, Matsunami) pre-coated with 0.01% poly-L-lysine solution (P8920, Sigma). A coverslip was gently placed over the sample, and the edges were sealed with VALAP (1:1:1 mixture of Vaseline, lanolin, and paraffin). Embryos were imaged using a spinning-disk confocal system (CSU-W1; Yokogawa Electric Corporation, Tokyo, Japan) mounted on an inverted microscope (Eclipse Ti-E, Nikon), equipped with a 100× objective (CFI SR HP Plan Apochromat Lambda S 100XC Sil, Nikon) and a sCMOS camera (Zyla 4.2, Andor). Imaging was performed at 21–22°C. Fluorescent imaging of GFP::Histone H2B and mCherry::PH was performed with 488 and 532 nm lasers, respectively, alongside differential interference contrast (DIC) microscopy. For live imaging of adult worms, adult worms were anesthetized in 1 mM levamisole in M9 buffer and transferred onto 2% agarose pad on a glass slide. After mounting, a coverslip was applied and sealed with VALAP. Imaging was performed using the same spinning-disk confocal system equipped with a 20× objective (CFI Plan Apochromat 20X, Nikon) and a sCMOS camera.

### Measurement of embryonic viability and egg yields

For counting laid eggs, individual young adult worm of the indicated strain was transferred to fresh NGM agar plate and incubated at 22°C for 24 hours. The total number of eggs (embryos and hatched L1 larvae) was counted to determine daily brood size. To assess embryonic viability, individual adult worms were dissected in M9 buffer on eight-well glass slides, mounted with a coverslip and sealed along the edges with VALAP, and incubated at 22°C for 48 hours. At least 15 worms per condition were analyzed, and data were pooled across replicates, and the percentage of arrested embryos was calculated. Statistical comparisons between diploid and tetraploid strains with the same genetic background were performed using Student’s *t*-test.

### Osmolality treatment

Adult worms subjected to *perm-1* (RNAi) were dissected in either undiluted egg salts (ES) buffer (118 mM NaCl, 40 mM KCl, 3.4 mM CaCl2, 3.4 mM MgCl2, 5 mM Hepes [4-(2-hydroxyethyl)-1-piperazineethanesulfonic acid], pH 7.2) or modified egg salts buffer (diluted or concentrated). The slide containing released embryos and dissected worm bodies were mounted with a coverslip on the eight-well glass slides without poly-L-lysine pre-coating, sealed with VALAP, and imaged as above.

Embryos were imaged using the above-mentioned spinning-disk confocal microscope setup. The DIC and fluorescent imaging were performed on embryos at 2–4 cell stages for over 30 minutes to evaluate abnormalities in cell cycle progression. Based on images, embryos were classified into four categories: cell division without any detectable delay or failures in chromosome segregation and cytokinesis as “normal division”, cell division with noticeable delay but without showing any division failures as “slow division”, cell division exhibiting errors in chromosome segregation or cytokinesis, or embryos containing abnormal shaped nuclei or multiple nuclei within an individual blastomere as “abnormal division”, and embryos showing no progression of cell cycle events as “arrested division”.

### Quantification of cellular parameters

For measuring the embryo size, the long (*L*) and short (*W*) axes of embryos at 2-cell stage were measured in the DIC images by using ImageJ software with a line ROI (region of interest) tool. For measuring the cell size, the cross-sectional area of AB blastomeres at prophase—identified by the presence of an intact nucleus with signals of GFP::histone H2B in the nucleoplasm just before the timing of nuclear envelope breakdown—was measured by aligning the elliptical section of ROI on the edge of blastomeres using ImageJ software. The plasma membrane of blastomeres were detected from DIC images, assisted by mCherry::PH signals in case using NF4496 and ECB003 strains. Cross-sectional areas of *perm-1(RNAi)* embryos under different osmolality conditions were normalized to mean values of untreated embryos in M9 buffer.

3D refractive-index (RI) imaging of embryos in M9 buffer was performed using a holotomographic microscope (HT-X1; Tomocube Inc) on an eight-well glass slide. Briefly, 2D holographic images were acquired from 48 azimuthally symmetric directions with specific polar angle controlled by a digital micromirror device. The 3D tomograms were reconstructed based on RI values in the equipped software. Mean RI values within individual embryos were calculated by defining ROIs inside each embryo (excluding boundaries) on Z-projected tomogram images using ImageJ software. Mass density was estimated from the equation, *n_(x,y,z)_ = n_m_ + αρ_(x,y,z)_*, where *n_(x,y,z)_* is the measured mean RI value within the embryo, *n_m_* is the RI value in the medium, *α* is the RI increment for protein and nucleic acid (0.190 mL/g; (Zhao *et al*., 2011)), and *ρ_(x,y,z)_* is the mass density (g/mL) (Biswas et al., 2025; Kim and Guck, 2020). Background RI values (i.e., the medium surrounding the embryos) were used as *n_m_* value.

For detection of intracellular biomolecules, embryos were prepared by the standard freeze-crack protocol on a glass-slide (Kimura and Onami, 2005). Embryos were fixed with −20°C methanol, and washed with Phosphate Buffer Saline (PBS; 136.8 mM NaCl, 8.04 mM Na_2_HPO_4_, 2.68 mM KCl, 1.763 mM KH_2_PO_4_, pH 7.4 [NaOH]). The fixed embryos were then stained for 15 minutes at 22°C with: 1 µg/mL BODIPY493/503 (D3922, Thermo Fisher Scientific) for neutral lipids, 5 µg/mL Cy5 NHS (N-hydroxysuccinimide)-ester (13020, Lumiprobe) for proteins, and 10 µM pyronin Y (14488, Cayman Chemical) for RNAs. Fluorescence imaging was performed using a wide-field microscope equipped with a 40× objective (CFI Plan Apochromat 40X, Nikon) and sCMOS camera. Mean fluorescence intensity within embryos was quantified using ImageJ software, subtracting background signals. The mean values in individual embryos from each condition and strain were normalized by the mean value obtained N2 strain without RNAi treatment. Statistical significance of RIs and fluorescence intensity differences was evaluated using the Wilcoxon signed-rank tests using the R software.

## ACKNOWLEDGEMENTS

We are grateful to Ririka Ikebe, Ayano Komachi, and other laboratory members for maintaining tetraploid strains and for the discussion. We also appreciate Dr. Kiyoji Nishiwaki at Kwansei Gakuin University for helping establish tetraploid worm strains. Some *C. elegans* strains were provided by the *Caenorhabditis* Genetic Center (CGC), which is funded by NIH Office of Research Infrastructure Programs (P40 OD010440). This study was supported by JSPS KAKENHI Grant Numbers JP20H03253 to Y.H.

