## supplemental figures for "Tetraploid *Caenorhabditis elegans* embryos exhibit enhanced tolerance to osmotic stress"

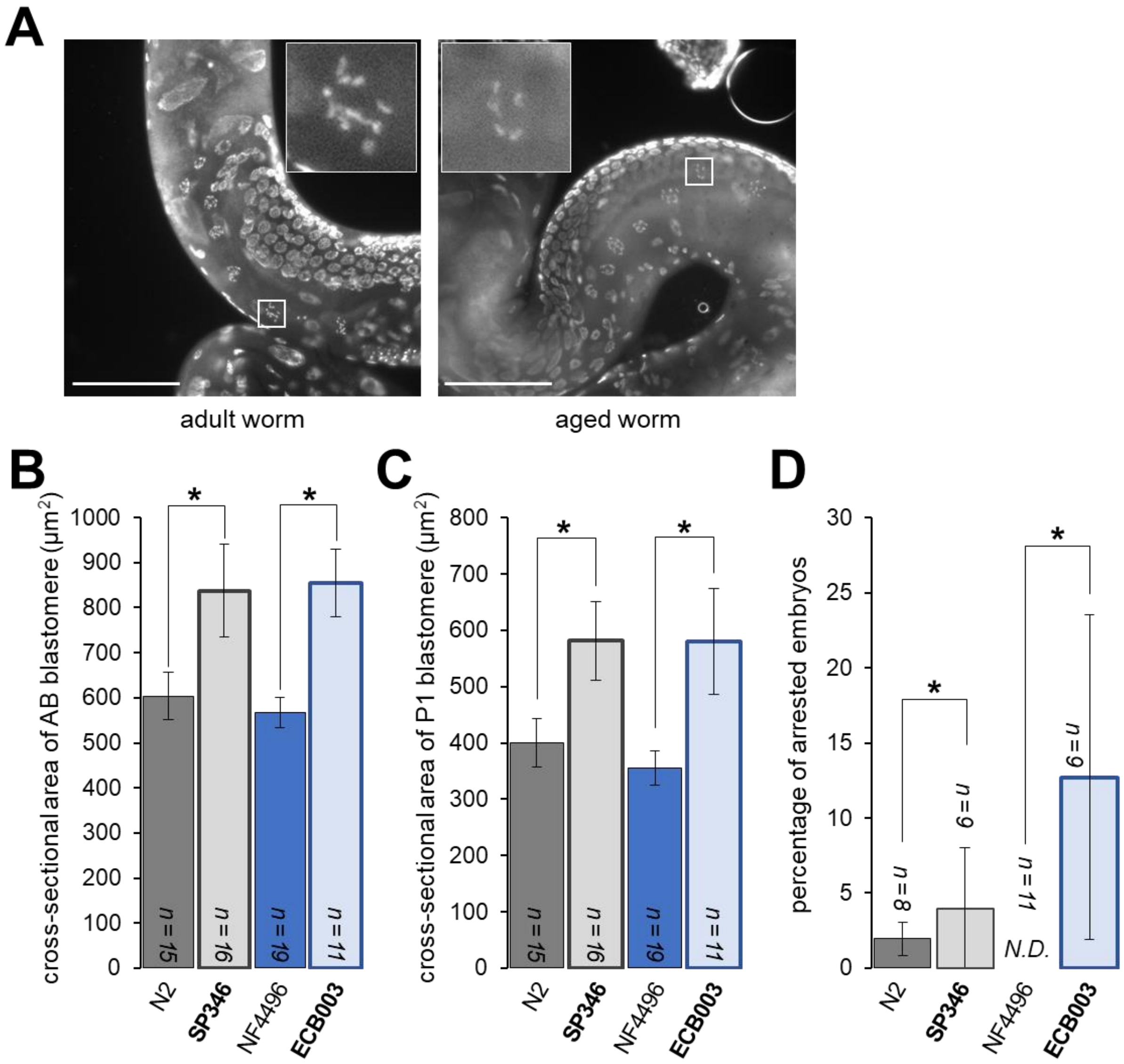


**Figure S1.** (**A**) Representative images of adult and aged tetraploid ECB003 worms stained with Hoechst 33342. The inset shows a magnified view of an oocyte region with visible chromosomes. Scale bar: 100 µm. (**B, C**) Mean cross-sectional areas numbers (± s.d.) of AB (**B**) and P1 blastomeres (**C**). (**D**) Percentage (± s.d.) of embryos that failed to hatch and arrested before reaching the larval stage (without hatching from eggshell) within 48 h of incubation on NGM plates. Asterisks indicate significant differences between tetraploid and diploid embryos (Student's t-test, $P < 0.005$). N.D. indicate the data not detected.

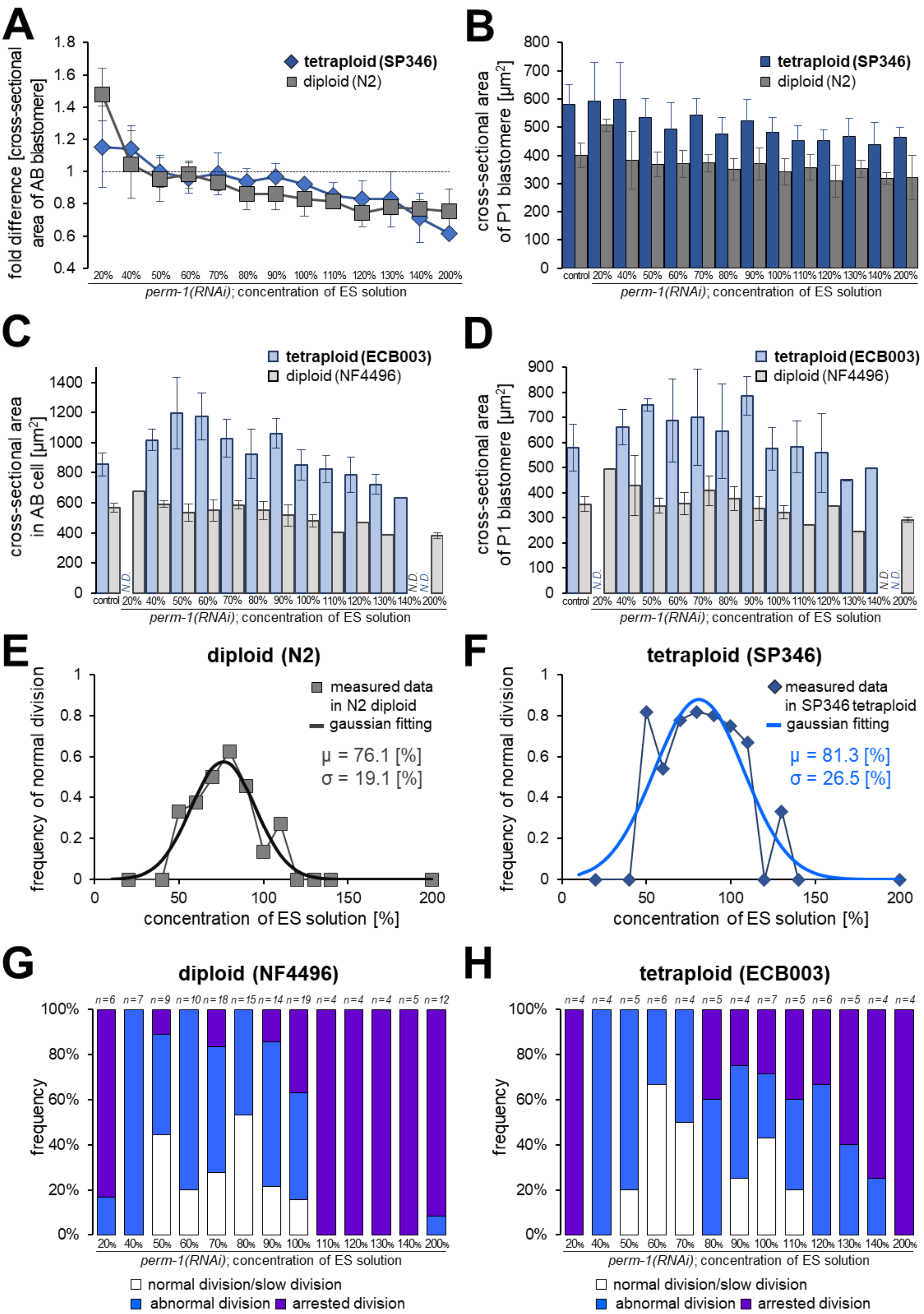
A
fold difference [cross-sectional area of AB blastomere]
tetraploid (SP346)
diploid (N2)
perm-1(RNAi); concentration of ES solution
B
cross-sectional area of P1 blastomere [μm²]
tetraploid (SP346)
diploid (N2)
perm-1(RNAi); concentration of ES solution
C
cross-sectional area in AB cell [μm²]
tetraploid (ECB003)
diploid (NF4496)
N.D.
perm-1(RNAi); concentration of ES solution
D
cross-sectional area of P1 blastomere [μm²]
tetraploid (ECB003)
diploid (NF4496)
N.D.
perm-1(RNAi); concentration of ES solution
E
diploid (N2)
frequency of normal division
measured data in N2 diploid
gaussian fitting
μ = 76.1 [%]
σ = 19.1 [%]
concentration of ES solution [%]
F
tetraploid (SP346)
frequency of normal division
measured data in SP346 tetraploid
gaussian fitting
μ = 81.3 [%]
σ = 26.5 [%]
concentration of ES solution [%]
G
diploid (NF4496)
n=6 n=7 n=9 n=10 n=18 n=15 n=14 n=19 n=4 n=4 n=4 n=5 n=12
frequency
perm-1(RNAi); concentration of ES solution
normal division/slow division
abnormal division
arrested division
H
tetraploid (ECB003)
n=4 n=4 n=5 n=6 n=4 n=5 n=4 n=7 n=5 n=6 n=5 n=4 n=4
frequency
perm-1(RNAi); concentration of ES solution
normal division/slow division
abnormal division
arrested division

**Figure S2.** (**A**) Fold changes in cross-sectional area of AB blastomeres in diploid (N2) and tetraploid (SP346) *perm-1(RNAi)* embryos in each ES concentration, normalized to untreated controls without *perm-1(RNAi)*. (**B**) Mean cross-sectional area (± s.d.) of P1 blastomeres in diploid (N2) and tetraploid (SP346) *perm-1(RNAi)* embryos incubated under different ES concentrations. (**C, D**) Mean cross-sectional area (± s.d.) of AB (**C**) and P1 (**D**) blastomeres in diploid (NF4496) and tetraploid (ECB003) *perm-1(RNAi)* embryos across concentrations. N.D. indicate no data were obtained. (**E, F**) Gaussian fits of the frequency of normal cell division across ES concentration in diploid (**E**, N2) and tetraploid (**F**, SP346) and *perm-1(RNAi)* embryos. Center ($\mu$) and width ($\sigma$) values of the fitting curve are indicated. (**G, H**) Frequencies of categorized cell division phenotypes in diploid (**G**, NF4496) and tetraploid (**H**, ECB003) *perm-1(RNAi)* embryos across ES concentrations. Numbers of embryos analyzed ($n$) are indicated.

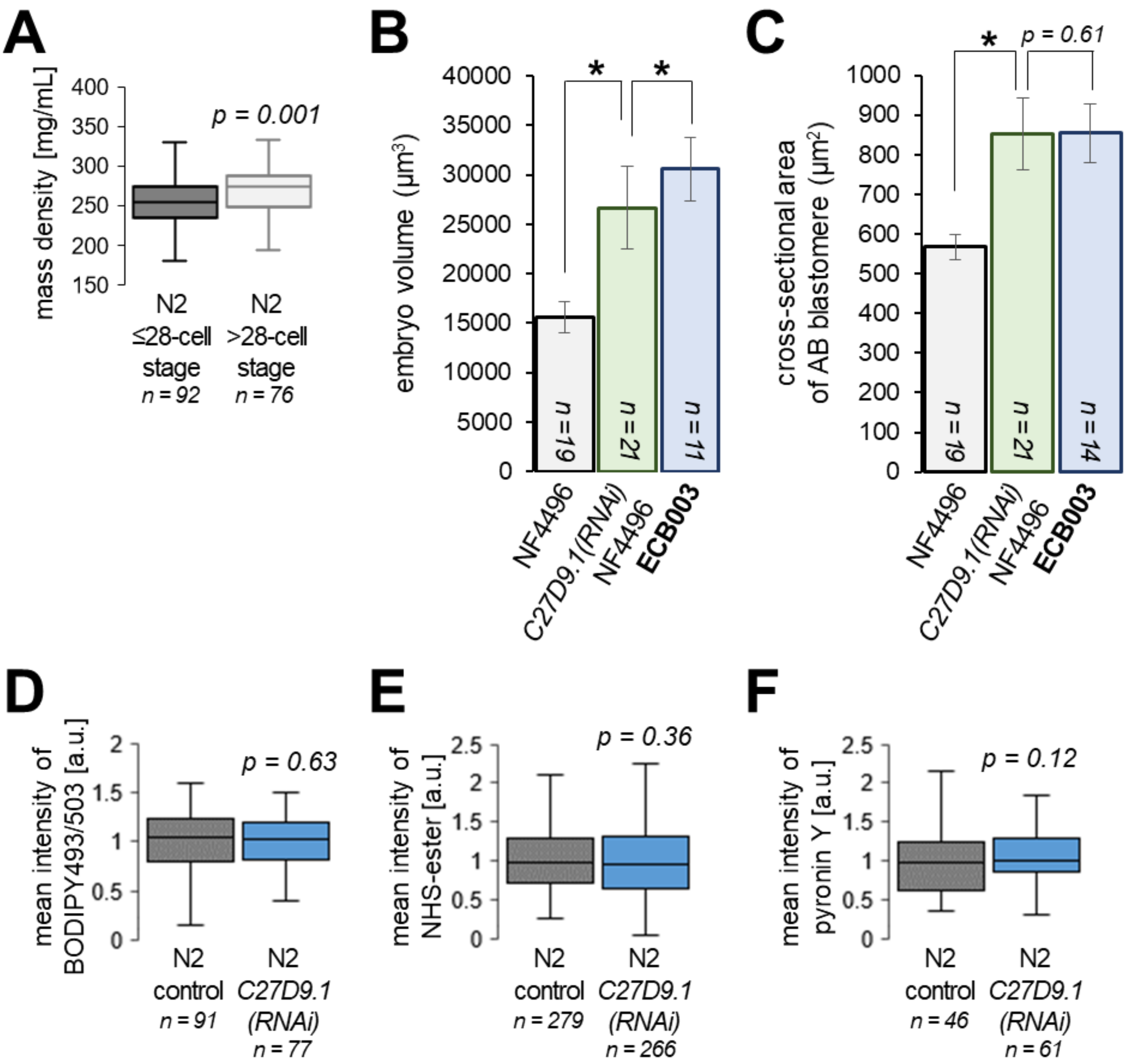


**Figure S3.** (**A**) Box plots of calculated cellular mass density derived from RI imaging in N2 embryos at less or more than 28-cell-stages. The line within the box represents the median. The upper/lower lines represent the 75th/25th percentiles. The whiskers represent the 95th/5th percentiles. *p*-value is from Wilcoxon tests comparing between them; numbers of embryos analyzed (*n*) are indicated. (**B, C**) Mean embryo volume (**B**, ± s.d.) and AB blastomere cross-sectional area (**C**, ± s.d.) at 2-cell stage. Asterisks indicate significant differences between data with different conditions (Wilcoxon test, $P < 0.005$). (**D–F**) Box plots showing normalized fluorescence intensities of BODIPY 493/503 (**D**), NHS-ester (**E**), and pyronin Y (**F**) in N2 embryos and *C27D9.1*(RNAi) embryos from N2. Intensities were normalized to mean N2 values.
